# Spatial Transcriptomics Sequencing of Mouse Liver at 2µm Resolution Using a Novel Spatial DNA Chip

**DOI:** 10.1101/2024.01.08.574734

**Authors:** Xun Ding, Kendall Hoff, Radha Swaminathan, Mikaela Koutrouli, Scott Pollom, Tianlong Huang, Xiaochi Li, Guoqiang Zhou, Zhicong Bai, Shizhe Yu, Zongping Xia, Lars Juhl Jensen, Filip Crnogorac, Su Yu, Glenn McGall, Jeremy Edwards, Wei Zhou

**Affiliations:** Centrillion Technologies, Palo Alto, CA 94303; Department of Biomedical Engineering, University of New Mexico, Albuquerque, NM 87131; Department of Chemistry and Chemical Biology, University of New Mexico, Albuquerque, NM 87131; Novo Nordisk Foundation Center for Protein Research, Faculty of Health and Medical Sciences, University of Copenhagen, Copenhagen, Denmark; Centrillion Technology, Hangzhou, Zhejiang Province, China 310052; Translational Medicine Center, The First Affiliated Hospital of Zhengzhou University, Zhengzhou, Henan, China 450052

## Abstract

Spatial transcriptomics enables analysis of gene expression that is spatially resolved within a tissue section, making it possible to elucidate the relationship between individual cells within the context of the tissue. This transformative technology enables researchers to better understand gene function within the context of health tissue, developmental processes, and disease. In this study, we present an innovative spatial transcriptomics technology and data using a high-resolution DNA chip with a total capture region size of 6.5 x 6.5 mm containing 2 x 2 µm features for spatial barcoding with no gaps between the features, thereby maximizing the capture area. These chips are manufactured at wafer scale using photolithography and are transferred to hydrogels, making them compatible with existing workflows for fresh frozen or paraffin-embedded samples. Herein, we examined a fresh frozen sample from an adult mouse liver. To analyze the data, we binned 10 x 10 features to represent a 20 µm x 20 µm capture area. We obtained 1.3 billion unique mapped reads, 68.78% sequencing saturation, with a median of 16,967 unique reads per region, indicating the potential for more unique reads with deeper sequencing. This high-resolution mapping of liver cell types and the visualization of gene expression patterns illustrate significant advancements in spatial sequencing technology.

## Introduction

The high-throughput analysis of spatially resolved gene expression at the single cell level, enabled by spatial transcriptomics technologies, has transformed our understanding of cellular processes and molecular mechanisms^1^. Traditional transcriptomics methods, such as bulk RNA sequencing, and newer single-cell methods provide invaluable insights into the gene expression patterns within a tissue or cell population; however, they overlook the intricate spatial organization of cells within complex biological tissues and organs^1,2^. The advent of spatial transcriptomics technologies has revolutionized our ability to capture gene expression data within its native spatial context, enabling us to understand tissue heterogeneity and interactions of cells within tissues both in healthy tissues and in models of development and disease^1,2^.

Spatial transcriptomics bridges the gap between high-resolution spatial information and transcriptomics data, allowing researchers to study the spatial distribution of gene expression across tissue sections^1–3^. This approach not only provides a detailed map of cellular activity within its anatomical context but also offers a unique opportunity to decipher the molecular signatures that underlie cellular heterogeneity^4^, tissue development^5,6^, and disease progression. By preserving the intricate relationships between neighboring cells and their surrounding microenvironment, spatial transcriptomics enables researchers to obtain enhanced biological understanding^3^, making it a pivotal tool in the fields of cancer research^4,7–11^, neuroscience^12^, developmental biology^5,6^, and drug development^13^. Molecular spatial analysis methods are divided into two approaches: imaging-based approaches and sequencing-based approaches. Imaging-based approaches suffer from low throughput, only capable of capturing the expression of a handful of transcripts per imaging run, whereas sequencing-based approaches suffer from low resolution^14^. This work demonstrates a significant improvement in the resolution of sequencing-based spatial transcriptomics methods.

Microarrays (or DNA chips) have been used in spatial transcriptomics. The DNA chips are manufactured with features or “spots” containing a poly-T capture sequence for mRNA capture and a unique barcode sequence to identify the spatial location of the feature to which the sequencing reads correspond. This barcode is sometimes called a zip code, since it encodes positional information. Due to limitations in the resolution of commercially available microarrays employed in spatial transcriptomics (10-55 µm)^15^, additional single-cell sequencing data has been combined with the spatial transcriptomics data to provide single-cell resolution. However, recently several new technologies have been introduced that provide improved spatial resolution but still introduce large gaps between features^16,17^.

Photolithographic microarray fabrication methods are currently used for the production of commercial microarrays with feature spacing of 4-10 µm^18,19^ We have previously demonstrated technological advances in wafer-scale manufacturing of DNA chips with the 5’ end tethered to a surface and a functional free 3’ end in hydrogels with sub-micrometer resolution^20^. Herein, our DNA chips were photolithographically manufactured at semiconductor scale using high-efficiency 5’-(2-nitronaphth-1-yl)benzyloxycarbonyl (“NNBOC”) phosphoramidite reagents with standard coupling, masking, alignment and exposure protocols^20,21^. Using these DNA chips, we demonstrate a spatial transcriptomics assay with fresh frozen adult mouse liver tissue sections.

The mammalian liver performs a diverse array of functions with approximately 50% of hepatocyte genes expressed in a spatially differentiated manner^22^. The liver is divided into lobules, repeating hexagonal units with approximately 12 concentric layers of hepatocytes^22,23^. Blood flows in from portal nodes and drains from central veins, creating a gradient of oxygen, glucose, and hormones^23^. Periportal hepatocytes function in a microenvironment of high oxygen and hormone levels and primarily function in gluconeogenesis, ureagenesis, cholesterol biosynthesis, protein secretion, and β-oxidation^22,23^. Conversely, the microenvironment of pericentral hepatocytes has high levels of Wnt (secreted by endothelial cells surrounding central veins^23^), low levels of oxygen, and low levels of circulating hormones^22,23^. The primary functions of these cells are glycolysis, bile acid production, glutamine synthesis, lipogenesis, and xenobiotic metabolism^22,23^. As an organ with such diverse and critical functions, the liver has been extensively studied using scRNA-seq and various spatial methods^22–25^. This makes the liver an excellent candidate for the evaluation of emerging spatial technologies.

## Results

High-resolution DNA chips for spatial transcriptomics were manufactured as previously described^20^. Briefly, semiconductor photolithographic manufacturing techniques were used in conjunction with highly-efficient light-directed DNA synthesis to prepare high-density DNA arrays on silicon wafer substrates. Positional information is encoded by the specific DNA sequences synthesized in a known spatial pattern on the silicon chips using photolithography^20^. High-resolution chips are fabricated by sequential exposures of lithography masks, each one directing a single base addition. The synthesized oligonucleotides are transferred from the silicon substrate to a thin polyacrylamide gel affixed to a glass slide while maintaining the spatial pattern, which inverts the oligonucleotides so they are anchored to the polyacrylamide gel at the 5’ end. The 3’ end of the spatially patterned oligonucleotides is free and able to participate in enzymatic reactions during the assay^20^.

The DNA chips can be manufactured at any size; for example, a 10 x 10 mm chip contains 5,000 x 5,000 (25 million total) 2 µm square features, and each wafer produces 111 10 x 10mm chips. Features are constructed adjacent to each other to cover the entire area without interstitial spaces (Fig. 1). Embedded within each feature is a unique predefined sequence, consisting of the partial Illumina sequencing primer 1 (used for PCR for amplification), a Unique Molecular Identifier (UMI), a zip code, and a 30 nucleotide poly-T sequence followed by 3’-VN-5’, which is optimized to capture the polyA region of an mRNA molecule just before the coding region (Fig. 1). The zip code encodes the X and Y coordinates that are decoded during bioinformatics analysis following sequencing (PostMaster™ software, Centrillion Tech, Palo Alto, CA). For the work presented herein, 6.5 x 6.5 mm synthesized chips were used.

**Figure 1:**
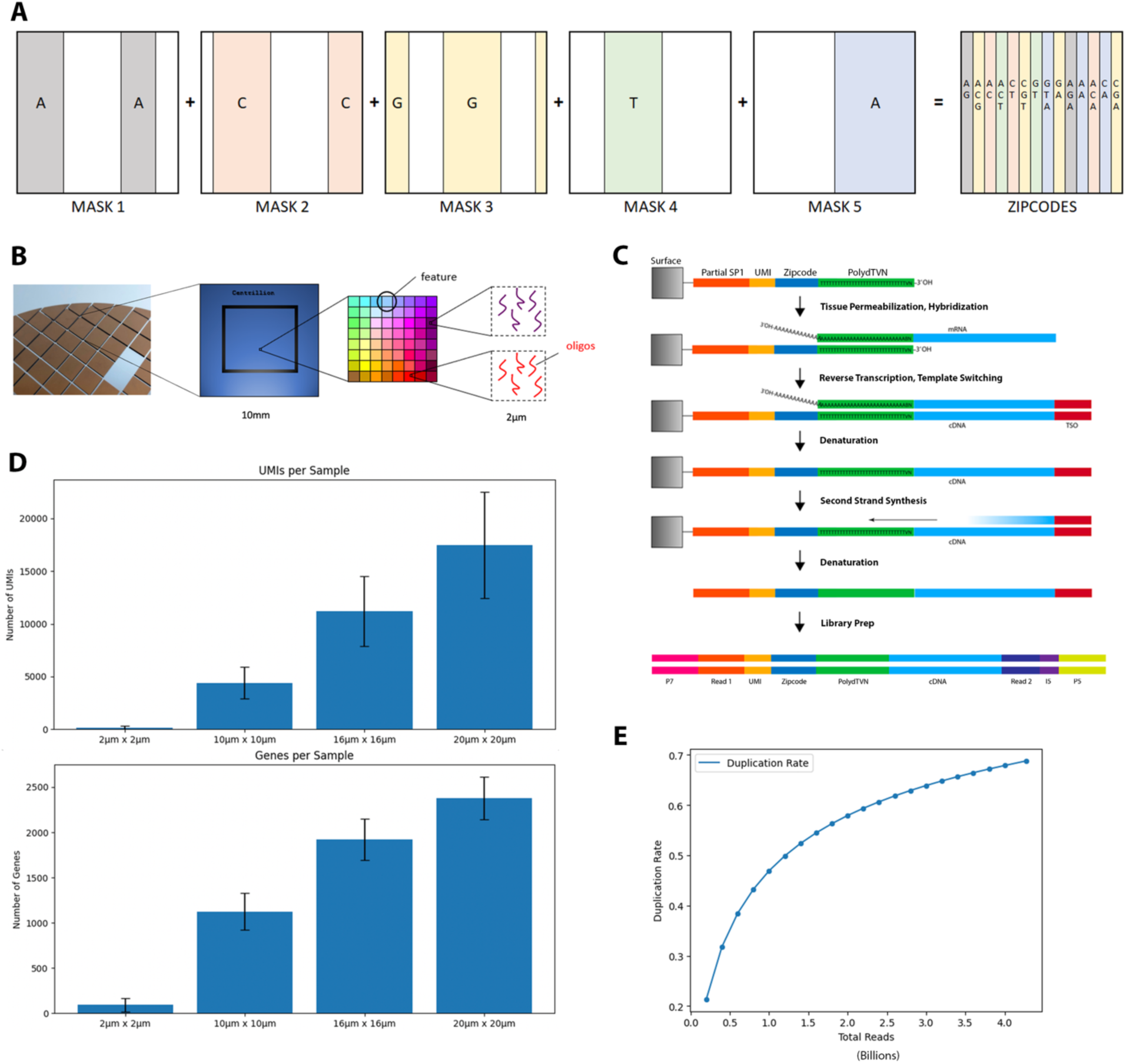
Chip Design and Performance. (A) Manufacturing of individual chips using photolithography. Vertical manufacturing of the zipcode design is shown, giving X positional information. Y positional information is achieved through the same methods with horizontal light-directed synthesis. **(B) Chip Design.** High-resolution spatial DNA chips are first manufactured using light-directed synthesis on silicon wafers then transferred to hydrogels as previously described^20^. Capture regions are confluent 2 x 2 µm squares. **(C) Assay workflow.** First, the sample is permeabilized and RNA from the tissue is hybridized to the chip. Chip oligos are extended and an adapter is added during reverse transcription with the use of a template-switching oligonucleotide (TSO). The mRNA is stripped away from the chip, and the chip sequence is copied during second strand synthesis. The second strand is then stripped from the chip and used in library preparation and downstream sequencing. The final sequencing library structure is shown. **(E) Chip performance.** Number of UMI and number of genes captured for binning sizes of 1, 5, 8, and 10. **(F) Sequence saturation.** Following deduplication, the saturation rate is calculated from the ratio of duplicated reads to the total number of reads— emphasizing the incremental accumulation of redundancy with the progression of sequencing depth.

Zip codes are designed using Gray Code to minimize errors during the synthesis of the DNA sequence^26^. The zip codes have a long-range minimum edit distance of 5, which indicates that if two zip codes have 5 or more differences, they are ≥ 5 units apart on the chip. Figure 1 illustrates the method of producing vertical zip codes on the array, which encode positional information along the X-axis. For horizontal zip codes encoding positional information along the Y-axis, the masks are rotated 90 degrees counterclockwise, and the process is repeated, creating a checkerboard pattern of unique X-Y zip codes. The zip codes are designed such that neighboring zip codes only differ by one base (Gray Codes), but their spatial extent covers the full chip area eliminating any unused border regions. Such a scheme allows for 100% surface coverage. In terms of resolution, high-resolution chips used in this work have individual features of unique zip codes that are 2 x 2 μm in size, however we have previously demonstrated this technology at 1μm resolution^20^. Notably, any individual mask contains polygons (exposure areas) much larger than the minimum feature size. Their overlay creates the minimum feature size, and, as such, the limiting factor is the overlay accuracy of the mask alignment equipment used (±0.3 μm at 3-sigma).

High-resolution spatial DNA chips are compatible with existing workflows for fresh frozen or paraffin-embedded samples. The data presented here uses a fresh frozen sample from mouse liver. Briefly, the tissue is fixed, decrosslinked, and permeabilized, and RNA from the tissue is hybridized to the chip (Figure 1). Following hybridization, RNA sequences are copied onto the chip using a reverse transcriptase and a template-switching oligonucleotide^27,28^. The original RNA strand is removed, and a second strand is synthesized, stripped from the chip, and used in downstream library preparation, which is finally sequenced using an Illumina system (Figure 1).

The spatial transcriptome library was sequenced in five separate runs using an Illumina Novaseq 6000. Each run generated read counts of 354,336,561, 501,709,129, 1,437,827,348, 1,596,786,045, and 3,099,518,653, respectively. This yielded a total of 6,990,177,736 reads. Of the totals reads, 6,194,279,449 reads contained a zip code. The reads with a decoded zip code were mapped to the mouse genome (GRCm39) using STAR^29^ and 4,271,034,555 reads were successfully mapped a unique location in the mouse genome, with 4,030,168,659 of these aligning to a specific gene using featureCounts^30^. Subsequent to this mapping, deduplication was performed. Reads that shared the same Ensembl gene ID, X position, Y position, and UMI were classified as duplicates; only the first occurrence was retained. Post-deduplication, a total of 1,333,303,108 unique reads remained. This implies a sequencing saturation rate of 66.9% of the library.

In order to understand sequencing depth and sensitivity, we analyzed the number of UMI and genes per feature at multiple bin sizes (summarized in Tables 1 and 2). Unbinned features (2 x 2 µm) generated between 16 and 3,396 unique reads per feature with a median count of 110, and between 1 and 976 different genes per feature with a median count of 70. At a binning of 10 (20 x 20 µm), corresponding to the approximate size of adult mice hepatocytes^31,32^, we generated between 8,002 and 79,375 unique reads per feature, and between 1,525 and 4,098 different genes per feature with a median count of 2,373. The capture efficiency observed is comparable to previously-reported scRNA-seq methods^33^.

**Table 1:** UMI captured at various bin sizes.

| Bin | Size | Mean | Median | Min | Max | IQR |
| --- | --- | --- | --- | --- | --- | --- |
| none | 2μm x 2μm | 168.66 | 110.00 | 16.00 | 3396.00 | 167.00 |
| 5 | 10μm x 10μm | 4390.92 | 4178.00 | 2001.00 | 28051.00 | 1457.00 |
| 8 | 16μm x 16μm | 11193.22 | 10869.00 | 5121.00 | 54340.00 | 2581.00 |
| 10 | 20μm x 20μm | 17467.38 | 16967.00 | 8002.00 | 79375.00 | 3917.50 |

**Table 2:** Number of genes captured at various bin sizes.

| Bin | Size | Mean | Median | Min | Max | IQR |
| --- | --- | --- | --- | --- | --- | --- |
| none | 2μm x 2μm | 93.05 | 70.00 | 1.00 | 976.00 | 89.00 |
| 5 | 10μm x 10μm | 1123.41 | 1107.00 | 395.00 | 2937.00 | 232.00 |
| 8 | 16μm x 16μm | 1921.97 | 1915.00 | 1041.00 | 3633.00 | 210.00 |
| 10 | 20μm x 20μm | 2377.98 | 2373.00 | 1525.00 | 4098.00 | 221.00 |

To evaluate the spatial accuracy in resolving spatial gene expression of the DNA chip and the sensitivity of our assay, we evaluated the spatial gene expression within the context of the tissue architecture and visualized the expression of specific cellular markers. The liver performs a vast array of metabolic functions that are regulated in spatial^22,23,34^, temporal^23,35^, and injury-related^34^ manners as determined by the microenvironment of each hepatocyte^22,23^. Zonation in the liver has been studied for over 80 years^36^. Liver lobules are hexagonally shaped with portal arteries and veins supplying blood at each of the 6 points of the hexagon and a central vein, through which blood is drained, at the center of the lobule^22,23,34^. Each of the concentric layers of hepatocytes has a microenvironment slightly different from its neighbors, defined by gradients of oxygen, glucose, and pancreatic hormones such as insulin, morphogens, etc^22,34^.

This structure enables production-line patterns of cells, as with the synthesis of bile salts, for example^23^. The first two enzymes in this cascade, CYP7A1 and HSD3B7 are expressed in the most pericentral cell layers where cholesterol, the starting material in the pathway, is most abundant^23^. The next-two enzymes in the cascade, CYP8B1 and CYP27A1, are expressed in the next layer, just periportal to those cells in layer 1^23^, indicating different steps of the enzymatic cascade occur in a spatially-regulated fashion. Researchers have further delineated the intricate zonation of opposing pathways in the liver including hepatic glucose metabolism (glycolysis and gluconeogenesis), glutamine synthesis and ureagenesis, lipogenesis and β-oxidation, protein secretion, and defense against pathogens and xenobiotics^22,23,34,37,38^. These distinct markers and intricate zonal distributions of cells emphasize the specialized roles of hepatocytes across these separate hepatic zones.

We began by examining the spatial expression of several known marker genes for various regions in the liver. The marker genes have been reported for cells at various hepatocyte layers as well as for other liver cells such as endothelial cells and Kupffer cells^22,23,34,35,39–41^. The dynamic expression patterns of several canonical marker genes can be observed in Figure 2. GLUL is expressed by pericentral hepatocytes^16,22,39,41^. It encodes glutamine synthetase, which converts glutamate into glutamine^40^. HAMP is expressed by midlobule hepatocytes^16,41^. It encodes hepcidin, which blocks the iron export protein, ferroportin, thus controlling iron storage in the liver and systemic iron availability^42–44^. CYP2F2 is a periportal hepatocyte markers^16,40,45^. There are 450 human cytochrome P450 proteins, which are involved in xenobiotic metabolism of which CYP2F2 encodes one^40,46^. From the expression patterns of these three markers, we can see well-defined tissue architecture that corresponds with tissue morphology observed in H&E images.

**Figure 2:**
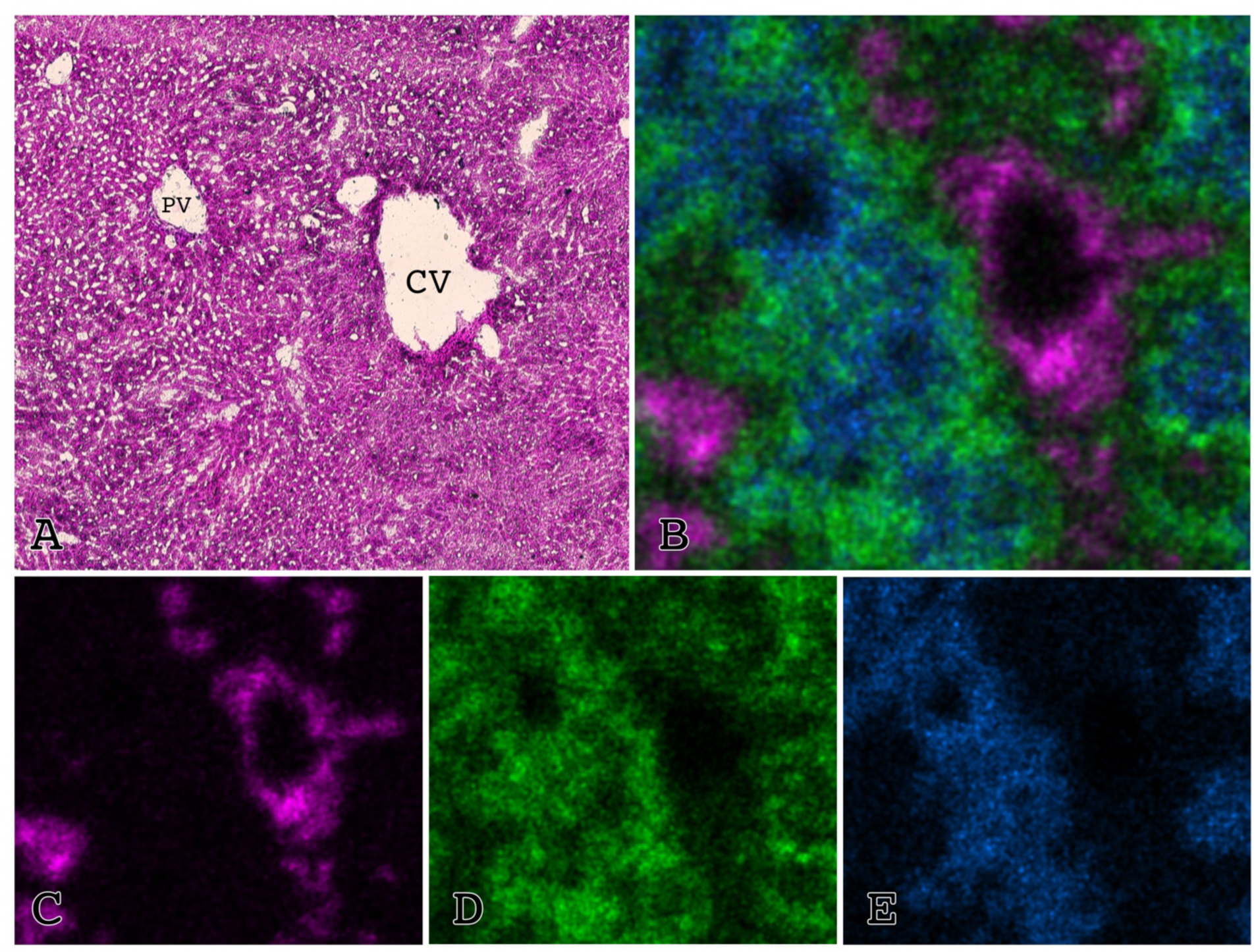
Spatial expression of cell-specific markers. **(A) H&E**. An H&E image from a neighboring section is shown with portal and central veins labeled. **(B) Merged.** Images for C-E are merged to show expression of pericentral (magenta), midlobule (green), and periportal (blue) hepatocytes markers. **(C) Pericentral Hepatocytes.** Expression of the pericentral hepatocyte marker, GLUL, is shown in magenta. **(D) Midlobule Hepatocytes.** Expression of the midlobule hepatocyte marker, HAMP, is shown in green. **(E) Periportal Hepatocytes.** Expression of the periportal hepatocyte marker, CYP2F2, is shown in blue.

Clustering methods were next applied to visualize the spatial composition of the tissue and identify the locations of the various cell types. Below, we will first apply Cell2location^47^ and the U-CIE^48^ tool, which translates high-dimensional data into colors, to generate supervised cell type spatial and cluster maps.

We applied the Cell2location spatial deconvolution method to interpret gene expression data within liver tissue sections, as referenced from the Liver Cell Atlas^49^. Cell2location was used to quantify the cell type composition at each tissue location, integrating this information to form a detailed cellular map. The analysis translated high-dimensional transcriptomic data into a resolved spatial context, revealing the distribution and abundance of cell types across the liver section. The result is a color-coded representation that provides insights into the tissue’s cellular architecture (Fig. 3), with the intensity of each color corresponding to the presence of specific cell types. This approach highlights the spatial heterogeneity of the liver and can be used to identify regions with distinct cellular functions or compositions, offering a window into the intricate structure and functionality of liver tissue at the cellular level.

**Figure 3:**
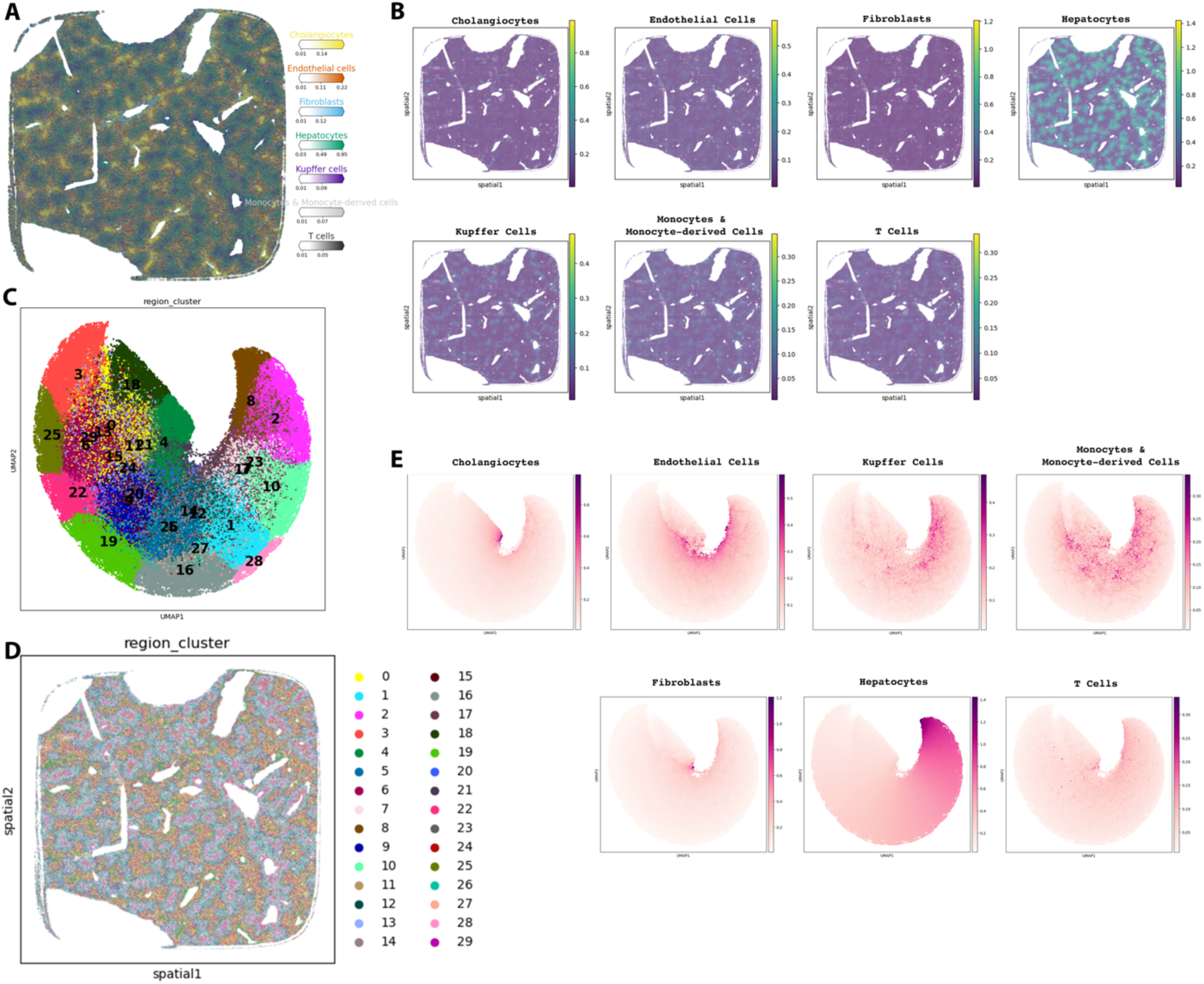
Cell type composition using Cell2Location. **(A)** Cell Type Composition Mapping via Cell2Location Deconvolution. A color-mapped image of a liver tissue section, indicating the cellular composition as determined by Cell2Location deconvolution. The colors represent the estimated proportions of different cell types within the tissue, with the color intensity corresponding to the prevalence of each cell type. Each color corresponds to a specific cell type as annotated in the legend, which quantifies the relative abundance of cholangiocytes, endothelial cells, fibroblasts, hepatocytes, Kupffer cells, monocytes & monocyte-derived cells, and T cells within the tissue matrix. The color gradients in the legend reflect the density of each cell type, providing a visual representation of the cellular heterogeneity across the tissue. **(B) Spatial Distribution of Cell Types in Liver Tissue Analyzed by Cell2Location and SCANPY/AnnData**. Each panel corresponds to a different cell type as indicated by the titles: cholangiocytes, endothelial cells, fibroblasts, hepatocytes, Kupffer cells, monocytes & monocyte-derived cells, and T cells. The color intensity within each panel reflects the relative abundance of the respective cell type across the tissue, with the color scale on the right indicating the density from low (purple) to high (yellow). **(C-E) Comprehensive Analysis of Liver Tissue Using Cell2Location Inferred UMAP Clusters. (C) Series of density plots for individual cell types including cholangiocytes, endothelial cells, hepatocytes, Kupffer cells, monocytes & monocyte-derived cells, and T cells.** These plots display the spatial distribution of cell-type-specific signals within the tissue section, shaded according to the intensity of the presence of each cell type. **(D)** UMAP visualization delineating regional clusters within the liver tissue, with discrete colors marking different clusters. Each number represents a unique region cluster identified by Cell2Location analysis, suggesting spatially distinct transcriptomic profiles. **(E)** Annotated liver tissue section with overlaid region clusters correlating to the UMAP analysis. The clusters are color-coded to match the UMAP plot, providing a spatial context for the cell type distributions inferred by Cell2Location, and highlighting the heterogeneity and complex structure of the liver tissue at the cellular level.

U-CIE is a software tool for translating high dimentional data into a color representation^48^. We used U-CIE to provide additional visual insight into the spatial pattern of gene expression in the mouse liver. U-CIE translates the spatial data into the CIELAB color space, therefore highlighting gene expression variations in a manner that can be interpreted visually. Essentially, U-CIE converts the high dimensional gene expression data at each liver location into a color, visually demonstrating the heterogeneity of the liver’s cellular landscape. Further, single-cell data from the Liver Cell Atlas^49^ is projected in the three UMAP dimensions of the spatial data, with U-CIE coloring applied to depict cell types. This combination reveals the distribution of cell types across the liver, providing a color-coded map of cellular organization (Fig. 4).

**Figure 4:**
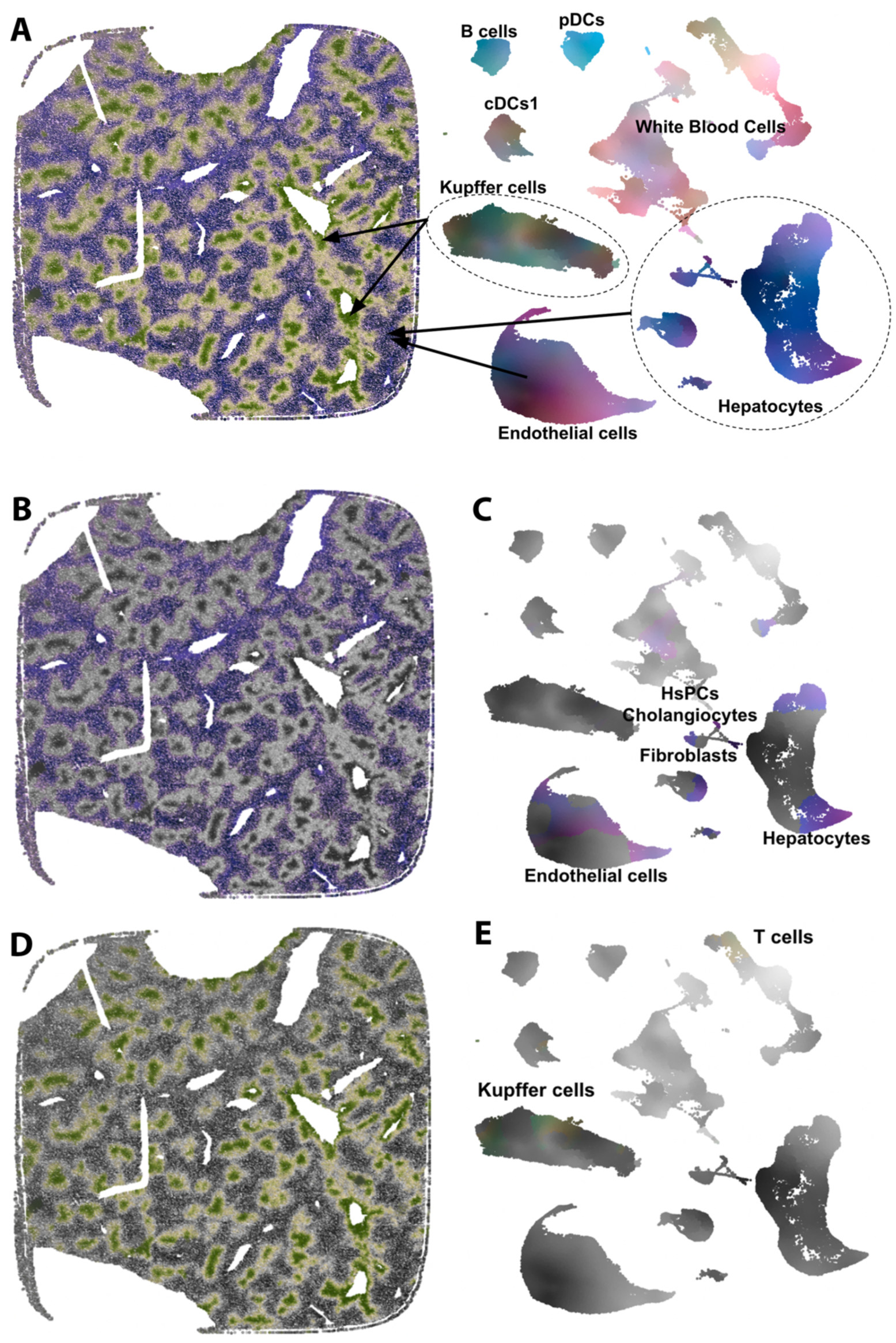
Integration of U-CIE and Liver Cell Atlas for Spatial Transcriptomics Analysis. **(A)** U-CIE color-coded liver tissue section, illustrating varying gene expression levels. Each image pixel corresponds to an individual transcriptomic profile, with distinct color patterns representing various gene expression signatures. UMAP visualization of single-cell data from the Liver Cell Atlas is displayed on the right. **(B-E)** Cell subtypes in the tissue, with each point signifying a single cell and the color-coding reflecting cell subtype annotations from the Liver Cell Atlas. Gray-scale images of the liver tissue are shown in **(B&D)**, with small, colored regions indicating cell type compositions based on Liver Cell Atlas annotation. The corresponding UMAP plots are shown in **(C&E)**, with the highlighted color indicating the cell type composition of the corresponding region in the tissue.

Additionally, unsupervised clustering was performed using Seurat v5^50^ to delineate subtypes of the observed cell types (Fig. 5). Unsupervised clustering enables us to identify patterns of expression in the tissue that may be missed with supervised clustering methods. Specifically, we identify several unique clusters of hepatocytes that vary in the expression of genes important to liver function. 17 total distinct clusters were identified using Seurat (Fig. 5), including three zones of hepatocytes as previously described^51^, and additional unique clusters within those zones which represent functional differences in cell layers. The result is eight concentric layers of hepatocytes within the lobule (Fig. 5D). These are periportal hepatocytes (cluster 0), pericentral hepatocytes (cluster 3), and 6 distinct midlobular hepatocytes (clusters 0-4, 6, 7, and 9), which is consistent with previous reports^35^. In order to further differentiate hepatocytes, we filtered for those features and reprocessed them using the Seurat pipeline (Fig. 5 E & F).

**Figure 5.**
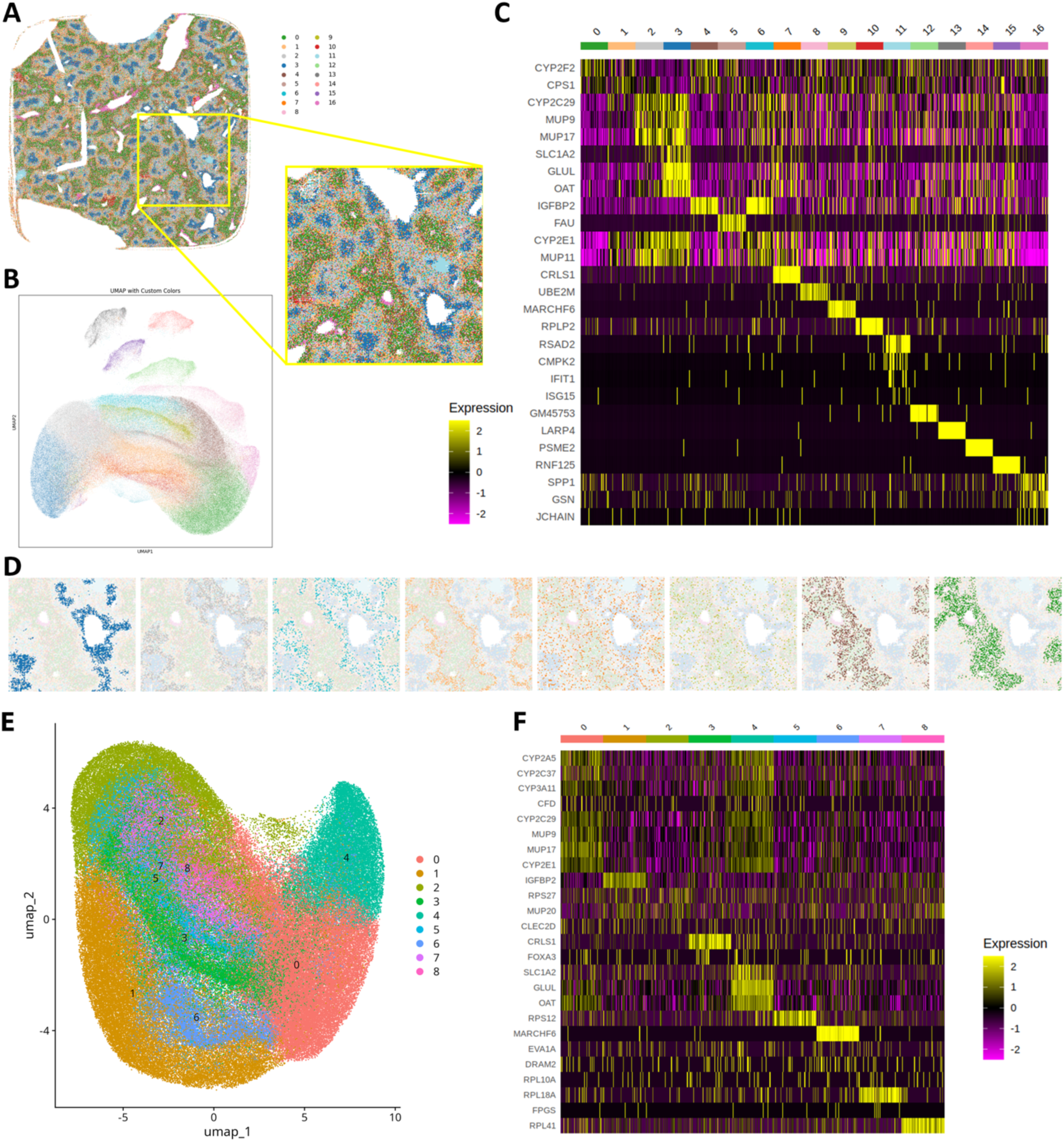
Successful capture of the mouse liver transcriptome. (A) Reconstructed map of spatial expression based on unsupervised clustering data. Hash marks represent barcode positions with each barcode being 0.2µm in height and width. (B) UMAP plot for clustering data. (C) Differential expression profile analysis of cell clusters. Seurat uses the expression of integrated most variable genes to assign PCA scores and determine clusters. Full expression levels are provided in Supplementary Materials. (D) Hepatocytes form 8 concentric clusters. Clusters are displayed from pericentral to periportal: clusters 3, 2, 6, 1, 7, 9, 4, and 0. (E) UMAP plot for clustering filtered for hepatocytes. The original clusters 3, 2, 6, 1, 7, 9, 4, and 0 were filtered for and clustering was repeated. We retain 8 unique clusters. (F) Differential expression profile for clustering filtered for hepatocytes. Top gene expression profiles for the new hepatocyte clusters. Full expression levels are provided in Supplementary Materials.

In addition to hepatocytes, clusters 5 and 10 were representative of B, T, and Kuffer cells; cluster 11 corresponded with erythroid cells and fibroblasts, and cluster 16 is comprised of plasma cells. Cluster annotation was performed manually based on expression profiles^52,53^. Many of the uniquely expressed genes seen in the clustering are important in disease, particularly hepatocellular carcinoma (HCC). Genes that were upregulated in unique clusters include IGFBP2, FAU, CRLS1, UBE2M, MARCHF6, RPL2, RSAD2, CMPK2, IFIT1, ISG15, LARP4, PSME2, RNF125, SPP1, GSN, and JCHAIN (Table 3). Changes in these genes and clusters associated with these genes will likely be important in future studies of liver disease.

**Table 3:** Genes uniquely expressed by individual. Genes listed show unique expression in distinct cell clusters as identified by unsupervised clustering with Seurat (Fig. 5). Many of these have been shown to be influential in disease; in particular, hepatocellular carcinoma. A brief description of their impact is listed.

| Cluster | Gene | Importance in Disease |
| --- | --- | --- |
| 4 and 6 | **IFGBP2** | IFGBP2 is a potent oncogene in HCC, driving epithelial-mesenchymal transition (EMT) through activation of the Wnt/β-catenin pathway in primary human tumors<sup>62</sup>. Overexpressed in HCC, drives EMT through Wnt/β-catenin pathway. |
| 5 | **FAU** | FAU is overexpressed in hepatic cellular carcinoma<sup>63</sup> and differentially expressed in between males and females with Hepatitis C Virus induced cirrhosis<sup>63,64</sup>. |
| 7 | **CRLS1** | CRLS1 has been implicated in many cancers, including HCC<sup>65</sup>. CRLS1 has been reported to be a tumor suppressor<sup>65</sup> with a positive correlation between high CRLS1 expression and survival<sup>66</sup>. Surprisingly, it has also been reported to confer resistance to ionizing radiation treatment in HCC<sup>67</sup>. |
| 8 | **UBE2M** | Drives HCC through negative regulation of p53<sup>68</sup> and increased cell proliferation via β-catenin/cyclin D1 signaling<sup>69</sup>. upregulation is associated with poor disease prognosis<sup>69</sup>. |
| 9 | **MARCHF6** | Promotes cell growth and migration through up-regulation of ATF2<sup>70</sup>. |
| 10 | **RPLP2** | Upregulated in several cancers, including HCC, and its overexpression is associated with poor prognosis<sup>71,72</sup>. RPLP2 activates TLR4 on the cell surface, which activates the PI3K/AKT/mTOR/HIF-1α pathway and |
|  |  | promotes cell proliferation, aerobic glycolysis, and cell growth<sup>71</sup>; it also inhibits ferroptosis<sup>72</sup>. |
| 11 | **RSAD2,**<br>**CMPK2,**<br>**IFIT1,**<br>**ISG15** | RSAD2 has been shown to play a protumorigenic role in HCC via increased lipogenesis and glycolysis<sup>73,74</sup>. Upregulation of viperin is inversely correlated with survival rates in patients<sup>73,74</sup>. CMPK2 has been shown to be upregulated in HCC cell lines<sup>75</sup>. ISG15 has been shown to trigger tumorigenesis and metastasis in HCC<sup>76,77</sup>, and it is a strong prognostic marker inversely correlated with 5-year survival in patients with HBV-related HCC<sup>78</sup>. |
| 13 | **LARP4** | LAPR4 is tumor suppressor in several cancers, affecting both cell growth and migration<sup>79</sup>. It has also been identified as a potential cause of HCC in sleeping beauty screens, although the mechanism is not well understood<sup>79</sup>. |
| 14 | **PSME2** | PSME2 serves as a biomarker of M1 macrophage infiltration in cancer<sup>80</sup>, but there is little research in HCC. When overexpressed in osteosarcoma, it suppresses tumor cell proliferation, migration, and invasion<sup>80</sup>. |
| 15 | **RNF125** | RNF125 is downregulated in HCC, which is associated with poor patient prognosis<sup>81</sup>. |
| 16 | **SPP1, GSN,**<br>**JCHAIN** | SPP1 has been shown to effect cell proliferation, immune invasion, and metastasis and cancer<sup>82</sup>. Increases in expression levels linked with increased immune cell infiltration and poor patient prognosis in HCC and other types of cancer<sup>82,82,83</sup>. GSN promotes EMT through enhanced cell |
|  |  | motility, and overexpression is correlated with poor prognosis<sup>84</sup>. JCHAIN is a prognostic marker for several types of cancer; favorable for head and neck and for breast cancer, but unfavorable in renal cancer<sup>52,53</sup>. |

## Discussion

This work demonstrates significant advances in the photolithographic manufacturing of spatial DNA chips with a highly optimized capture surface. High-resolution spatial DNA chips were manufactured at semi-conductor scale using photolithography. This production technique is suitable for large-scale and highly efficient manufacturing. The development of high-resolution chips represents a pivotal advancement in spatial transcriptomic technologies, merging positional information with extensive tissue coverage and efficient mRNA capture. To achieve these results, we utilized a combination of chemistry, chip design, and bioinformatics development.

The mammalian liver is a well-characterized, complex tissue containing many different cell types that perform coordinated metabolic functions. This makes it an excellent tissue to demonstrate the capabilities of new spatial sequencing technologies. We were able to validate the use of the chips with mouse liver, achieving sequencing results comparable to previously reported scRNA-seq data. This underscores the chip’s potential in comprehensively understanding tissue architectures and in early-stage pathological detection.

At 69% sequencing saturation, our sample yielded over 643 million effective reads from a single section with a median of 12,967 UMIs and 2,373 genes per 20 x 20 µm area (the average size of an adult mouse liver cell), demonstrating exceptional molecular capture capacity. By integrating the analysis techniques of U-CIE and Cell2Location, we have been able to decode complex gene expression patterns within liver tissues, enhancing our understanding of cellular diversity and structure. We believe the advancements demonstrated here establish a new benchmark for future research in the field. As the field evolves, integrating computational strategies and advanced microscopic methodologies with platforms like this will be paramount to deepening our insights into tissue dynamics and cellular intricacies.

Despite the advancements associated with the high-resolution chip, challenges persist. Delineating precise cell boundaries and achieving accurate single-cell genetic mapping remain areas of active research^54^. Potential solutions may involve the deployment of advanced algorithms and possibly integrating machine learning techniques to enhance spatial transcriptomics data interpretation^47,55,56^. Additionally, utilizing markers from specific subcellular compartments, such as pre-mRNA, and merging high-resolution microscopy might further refine cellular boundary determinations in tissues with intricate architectures^55^.

## Materials and Methods

### Chip Fabrication and Decoding

The high-resolution chips, fabricated by Centrillion Technologies, are synthesized on 150mm silicon wafers using aligner model CENAL1000 (Centrillion Taiwan), generating 111 10mm x 10mm chips per wafer. Chips are singulated and subsequently inverted and transferred to a thin polyacrylamide gel (10-20 µm) cast on a silanized glass slide. This process has been previously described^20^. Each of the 10 x 10 mm chips contains 5,000 x 5,000 features that are 2µm x 2µm each and juxtaposed adjacently without interstitial spaces. Embedded within each feature is a predefined sequence, consisting of a fragment of the Illumina read primer 1 for amplification, a Unique Molecular Identifier (UMI), a zip code, and a 3’ sequence, 3’-TTTTTTTTTTTTTTTTTTTTTTTTTTTTTTVN-5’, which is optimized to capture the polyadenylated region of an mRNA molecule closest to the coding region. The zip code is constructed from a 15-16 nucleoside sequence indicative of the X coordinate, a ’GGG’ separator, and another 15-16 nucleoside sequence corresponding to the Y coordinate. This zip code can be effectively decoded using the PostMaster software.

The sequences common to all barcode probes (poly-T, UMI & primer), which required no spatial definition, are synthesized using standard 5’-DMT deoxynucleoside synthesis reagents & coupling protocols^20,21^. The spatially-defined zip code sequences are synthesized photolithographically using high-efficiency 5’-(2-nitronaphth-1-yl)benzyloxycarbonyl (“NNBOC”) phosphoramidite reagents with standard coupling, masking, alignment and exposure protocols^20,21^. Vacuum contact lithography was performed using chrome on quartz masks. Two-micron features are achieved by employing a modified masking protocol as described in Figure 1.

Spatial barcodes (aka zip codes) are designed such that the “embeddings” of adjacent barcodes have an edit distance of 1. The term “embedding” refers to the binary sequence that specifies when the barcode’s position on the chip is exposed to light during the chip synthesis via photolithography^26^. The embeddings of all the chip barcodes form a two-dimensional Gray Code^26^. Another important property of pairs of DNA barcodes on the chip is the minimum edit distance between barcodes that are sufficiently far apart. DNA barcodes that are 1, 2, 3, or 4 units apart have an edit distance equal to the number of units apart, whereas DNA barcodes that are at least 5 units apart have an edit distance of at least 5. This property holds true for both sections of the DNA barcodes, meaning that if the X and Y coordinates of a pair of barcodes each differ by at least 5, the edit distance of the first sections will be at least 5, and the edit distance of the second sections will be at least 5. The combination of using gray codes and maintaining a minimum edit distance for distant DNA barcodes enables the chip to be fully covered by DNA barcodes, ensuring efficient transcript capture, and allowing for error correction during both chip synthesis and subsequent decoding.

### Bioinformatics

The bioinformatics pipeline is outlined in Figure 6. The spatial sequencing data processing pipeline was constructed utilizing binary decoding software, PostMaster, developed by Centrillion Technologies, Palo Alto. PostMaster parses Unique Molecular Identifiers (UMIs), and X, Y coordinates from the raw fastq files. It then decodes these coordinates, comparing them to the theoretical sequence of the decoded zip codes. This decoding information, including edit distances and UMI, is stored within the sequence identifiers of a new R2.fastq.gz file. Only coordinates with an edit distance less than or equal to three (compared to the set of theoretical sequences) are retained for further analysis. 97.36% of reads were decoded for the data presented. Within this decoded subset, 1.20% of X zip codes and 1.30% of Y zip codes had an edit distance equal to exactly 3. Following decoding, sequences were trimed with Cutadapt^57^.

**Figure 6:**
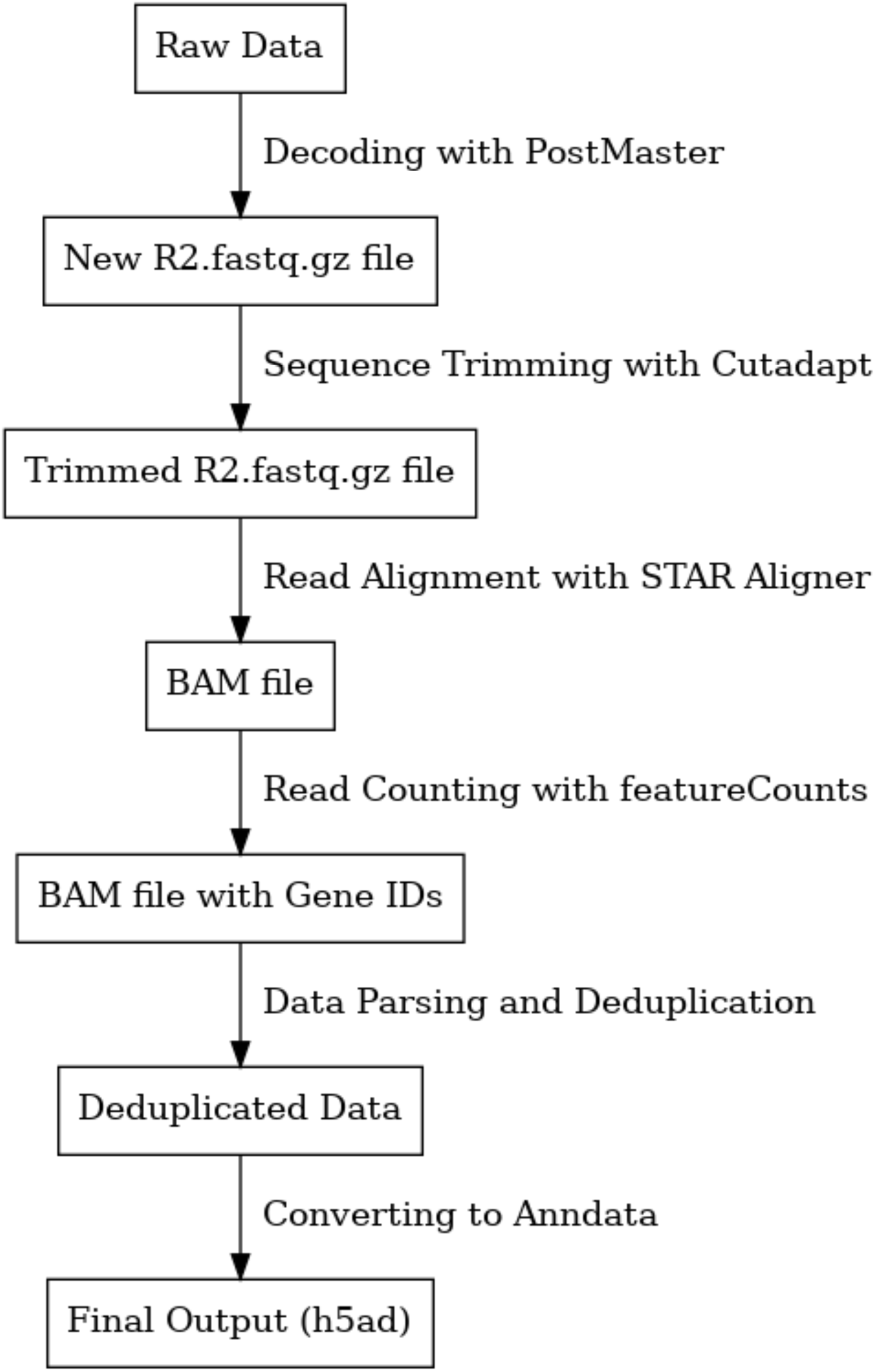
The bioinformatics pipeline. This study implemented a custom bioinformatics pipeline for the processing of spatial transcriptomics data obtained from paired-end FASTQ files. Our method primarily consisted of five key steps: 1. Sequence Parsing and Decoding with PostMaster: Our initial preprocessing step involved the utilization of PostMaster software. This tool was specifically designed to parse sequencing data from read 1 sequence, extract unique molecular identifiers (UMIs), and decode x and y zipcodes. PostMaster then provides the x and y coordinates as well as the edit distance of each zipcode relative to the theoretical sequences. The output of this step is a new R2.fastq.gz file with decoding information stored in the sequence ID (SeqID). 2. Sequence Trimming with seqtk: Following decoding, we applied the Cutadapt^57^ tool to trim the sequences in the newly formed R2.fastq.gz file. This process is vital in the removal of sequencing adapters and other potential sequence contaminants. 3. Read Alignment with STAR Aligner: Subsequent to trimming, we aligned the cleaned sequences to a reference genome using the Spliced Transcripts Alignment to a Reference (STAR) software. The alignment parameters were set to ensure only unique matches to the reference genome were retained, thereby eliminating multi-mapping reads. This step resulted in a Binary Alignment Map (BAM) file. 4. Read Counting with featureCounts: After alignment, we employed the featureCounts tool to perform a gene-level quantification. Specifically, this step involved counting the number of reads that were successfully mapped to each gene in the reference genome. FeatureCounts was executed three times using distinct annotation files to identify the mapping of reads to specific genomic regions: the last 200 bases of the 3’ end of a transcript, other regions within an exon, or introns. For subsequent analyses such as gene plotting, clustering, U-CIE, and Cell2Location, the data were pre-filtered. Reads that mapped to the last 200 bases of the 3’ end of a transcript were specifically included for analysis. 5. Data Parsing with bamreader.sh: After featureCounts, we utilized a custom script, bamreader.sh, to parse the decoding information embedded within the sequence IDs in the BAM file. The output of this step was a CSV file containing detailed information such as gene_id, x and y coordinates, edit distance, and UMI, which can then be used for subsequent downstream analyses. 6. Converting CSV file to Anndata: The final step of the data processing pipeline converts the CSV output into a h5ad file, which is a binary format used to efficiently store large, multi-dimensional arrays. This is accomplished through a custom Python script using the Scanpy library, named h5ad.py. In this step, any duplicate gene expression data, based on Unique Molecular Identifiers (UMI), X-Pos, Y-Pos, and Gene-id, are removed to ensure only unique spatial gene expression information is retained. The script then transforms this data into a sparse matrix representation, which helps to efficiently store and process the large amount of zero entries typical of gene expression data. Each unique gene and its corresponding spatial coordinates are recorded in this matrix, which is organized by gene symbols. Finally, an AnnData object is created with the sparse matrix, enabling the integration of gene expression and spatial data. This h5ad file provides a convenient and efficient format for subsequent data analysis tasks.

Subsequently, the trimmed reads are mapped to the reference genome using the STAR aligner^29^. These mapped reads are then processed by FeatureCounts^30^ to assign them to genomic features, thus generating a count matrix. Our custom script, bamreader.sh, utilizes SAMtools^58^ to read the .bam output from featureCounts, and parses the decoding information stored in the sequence identifiers, creating a CSV file with gene ID, UMI, and spatial coordinates. Bamreader.sh

Finally, an in-house Python script converts this data into an h5ad file, a format suitable for spatial transcriptomic analysis. This script also removes duplicates, ensuring the output consists of unique spatial gene expression data. Reads are considered ’similar’ if they have identical X-Pos, Y-Pos, Gene-ID, annotation regions, and chromosome number, and if the absolute differences in their start and end coordinates and mapping lengths on the reference genome are less than 2. Additionally, their UMIs must have an edit distance of 1 or less. Similar reads are grouped into clusters using a union-find algorithm, which aggregates reads into the same group even if not all members are directly similar to each other. After deduplication, only one representative read from each cluster is retained. The script constructs an anndata object, effectively integrating the gene expression and spatial data and as stored as an h5ad file^59^.

### Spatial Library Construction

The spatial transcriptomics procedure is initiated with tissue fixation, permeabilization using pepsin, and decrosslinking. mRNA was captured through hybridization then copied using reverse transcription with a template-switching oligo. Templating RNA molecules were removed and the cDNA was copied using second-strand synthesis. The second strand was denatured from the chip and used in downstream library preparation. The library was subsequently purified and size-selected.

### Pre-processing of Tissue

Tissue slides, retrieved from a -80℃ freezer, were briefly warmed at 37℃ for 1 minute. Slides were fixed using 100μl of 4% formaldehyde for 30 minutes at room temperature. Subsequently, decrosslinking was performed at 70℃ over 20 minutes in TE Buffer (10mM Tris-HC1;1mM EDTA). The slides were then swiftly rinsed in a 1×PBS-filled staining jar. For permeabilization, after removing excess water, the slides were treated with 100μL of 1/1000 dilution of 0.1% Pepsin and incubated at 37℃ for 5 minutes.

### Hybridization

For hybridization, a 20μL solution was formulated on ice with 9.4μL nuclease-free water, 5 μL 20× SSC, 2.4 μL 25mM MgCl2, 0.2μL 0.1M DTT, 2μL 10mM dNTPs, and 1μL RNase Inhibitor A (NEB, M0314L). A new Sequoia 2 chip was retrieved from the original container. After drying the surrounding of the chip’s gel area, any residue liquid was quickly aspirated. Both the chip gel and the designated tissue area on the slide received 10μL of Hybridization Buffer. The slide was subsequently inverted onto the chip, ensuring a snug, bubble-free contact. This configuration was incubated at 42℃ for 4 hours in a humidity chamber.

### Reverse Transcription

For reverse transcription on the chip, a 40 μL RT MIX was prepared on ice, containing 4μL PEG8000, 0.4μL Triton X, 1.5μL TSO Oligo with sequence 5‘-AAGCAGTGGTATCAACGCAGAGTACATrGrGrG, 2μL dNTPs, 1.6μL Tris HCl (pH 8.3), 1.2μL NaCl, 4μL MgCl2, 0.53μL GTP, 3.2μL DTT, 1μL RNase inhibitor (NEB, M0314L), 0.8μL Maxima H RT Enzyme (Thermo Fisher, EP0751), and 19.77μL nuclease free Water. After hybridization, the chip was rinsed in a staining dish with 0.1×SSC until the tissue slide released. A frame-seal (Biorad, SLF0201) was then aligned with the dashed region of the chip gel. The holes of the frame were filled with the RT Mix. The prepared chip underwent incubation in a pre-heated preservation box at 42℃ for 90 minutes and subsequently at 53℃ for 30 minutes.

### Second Strand Synthesis

For second strand synthesis, a 50μL reaction mix was prepared on ice, using 24μL nuclease-free water, 5μL 10X Thermopol Buffer, 4μL Bst Pol Full Length (NEB, M0328S), 7μL 10mM dNTPs, and 10μL of a primer with the sequence AAGCAGTGGTATCAACGCAGAG. Wash Buffer SSB was prepared using 13.2μL nuclease-free water, 1μL 10X Thermopol Buffer, 2.8μL 10mM dNTPs, and 4μL primer. The chip was sequentially cleaned: three times using 75μL nuclease-free water, incubated for 2 minutes; twice with 75μL 0.08M KOH for 5 minutes; and twice with 75μL Tris pH 7.5 for 2 minutes. Chips were washed in 20μL Wash Buffer SSB and then 50μL Second Strand Synthesis Mix was applied. The chip underwent a 30-minute incubation in a 65°C humidity chamber. The chip was then rinsed twice with 100μL Buffer EB and stripped with 20μL 0.08M KOH for 10 minutes. After mixing, 20μL was transfered to a 0.2ml PCR tube. This stripping process was repeated with 20μL 0.08M KOH for 10 minutes, and the second eluate was combined with the first. The pH was neutralized using 5μL of 1M Tris pH 7.0.

### cDNA Amplification and Purification

For cDNA pre-amplification, a mixture consisting of 50μL KAPA HiFi Hot Start Ready Mix (Roche, KK2601), 35μL cDNA, and 15μL of primers (Primer-f: CTACACGACGCTCTTCCGATCT; Primer-r: AAGCAGTGGTATCAACGCAGAG) was prepared in a total volume of 100μL. The PCR regimen included: initial denaturation at 98℃ for 3 minutes, followed by 14 cycles at 98℃ for 15 seconds, 63°C for 20 seconds, and 72°C for 1 minute, concluding with a 5-minute final extension at 72 °C. Purification of cDNA was performed using 0.7X Clean NGS-R Beads (CleanNA, CNGS-0050The final elution yielded 20μL of size-selected cDNA, which was transferred to a fresh PCR tube for further analysis.

### Fragmentation, End Repair, A-tailing and Ligation

For fragmentation, end repair, and A-tailing, an FS DNA Mix was prepared, consisting of 20μL Input DNA, 10μL Smearase® Mix (Yeasen, 12619ES24), and 30μL ddH2O. After gentle mixing and brief centrifugation, the mixture was subjected to a PCR program with a lid temperature set at 85℃. The regimen began with fragmentation at 30℃ for 2 minutes, followed by end repair and A-tailing at 72℃ for 30 minutes, and then held at 4℃.A Ligation Mix consisting of 60μL dA-tailed DNA, 20μL 5×Novel Ligation Buffer, 5μL Novel T4 DNA Ligase (Yeasen, 12626ES24), 3μL DNA Adapter (6μM, annealed from adapter_A (/5’Phosph/GATCGGAAGAGCACACGTCTGAACTCCAGTCAC) and adapter_B (5’-GCTCTTCCGATCT)), and 12μL ddH2O was incubated at 25°C for 15 minutes.

Following ligation, the products underwent purification with Clean NGS-R Beads. Briefly, beads were mixed with the product, washed, and the DNA eluted. A size selection targeting 300-700 bp fragments was achieved using a stepwise application of 0.6× and then 0.8× bead volumes. The final purified DNA was resuspended, with 25μL of the resultant solution transferred to a new PCR tube for subsequent analyses.

### Amplification and Adapter Addition

For library construction, a 50μL reaction mix was prepared using 25μL KAPA HiFi Hot Start Ready Mix, 2.5μL NEBNext Index primer (NEB, E7335S), 2.5μL NEB Universal Primer, and 20μL of purified ligated product. The PCR amplification was executed with an initial denaturation at 98°C for 45 seconds, followed by 14 cycles of 98°C for 20 seconds, 67°C for 30 seconds, and 72°C for 20 seconds. This was finalized with a 72°C extension for 1 minute and then held at 4°C indefinitely. Post-amplification, the library underwent size selection using Clean NGS-R Beads. Briefly, beads (0.6× and 0.8× volumes) were applied in successive steps, with washes and magnetic separation. The purified library was eluted in 30.5μL nuclease-free water, with 30μL transferred to a new PCR tube.

### Animal Husbandry and tissue preparation

Male C57BL/6 mice, eight weeks postnatal (PN), were used in this study. All procedures involving animals were conducted in strict accordance with the ethical guidelines established by the Institutional Animal Care and Use Committee of Zhengzhou University and were in compliance with the current laws regarding animal use in scientific research in China. Following perfusion, liver was harvested, embedded in OCT, stored at -80°C, and sectioned at a thickness of 10µm.

### Hematoxylin and Eosin (H&E) Staining

A section 20µm away from the section used for spatial transcriptomics was selected for H&E staining. Using the Beyotime C0105S Hematoxylin and Eosin Staining Kit, tissue samples were fixed with 4% paraformaldehyde at room temperature for 10 minutes. They were then stained with hematoxylin for 5 minutes, rinsed in water for 2 minutes, differentiated in 0.5% hydrochloric acid ethanol for 1 minute, followed by a 1-minute soak in 1xPBS. Finally, eosin staining was applied for 1 minute, and excess dye was removed with a 2-minute water rinse.

### Computational analysis: Data binning

Cho et al. showed that 10µm sided grids produced a much less noisy UMAP relative to 5µm square grids and was simultaneously able to discover clusters with single cell resolution^16^. Therefore, we combined counts of 5 consecutive features, performed convolution using Gaussian kernel with a width of 1 feature resulting in 10µm-sided binned features.

### Gene expression plots

Gene expression for genes of interest was plotted in greyscale using X and Y positional information for each transcript and a custom Python script. Black represents zero expression and white represents the highest level of expression. We used a bin size of 5, corresponding to 10 x 10 µm regions. This is smaller than typical adult mice hepatocytes, which are approximately 20 x 20 µm^31,32^. Pseudocolors were added and a merged image was generated by overlaying the colored images.

### Identification of regions within mouse liver tissue using unsupervised clustering

A threshold for total transcript count was applied to include only those binned features that were overlaid by the tissue. This binned count matrix was processed using Seurat v5 R package to identify regions present within the tissue. Briefly, the binned count matrix was normalized using Seurat’s SCTransform function. Clusters using the normalized binned count matrix were identified using Seurat’s shared nearest neighbor graph-based algorithm implemented in the FindNeighbors and FindClusters functions. The clustering was performed using an SLM algorithm with a resolution of 0.5.

### Visualization and annotation of clusters

A custom python script was used to visualize clusters in the context of the tissue section. Marker genes for each cluster were identified using Seurat’s FindAllMarkers function with the following parameters: min.pct = 0.25, logfc.threshold = 0.25.. The marker genes identified in this study were compared to other studies to annotate clusters generated by unsupervised clustering.

### Spatial Transcriptomics Data Analysis Using Cell2location

The analysis was performed using the Cell2location library, specifically employed for spatial transcriptomics data deconvolution. The spatial transcriptomics data were loaded using SCANPY^60^, targeting the H5AD file that contains the preprocessed spatial gene expression matrix. Similarly, annotated single-cell RNA-seq data from the Liver Cell Atlas was loaded as a reference dataset. Preliminary examination and preprocessing were performed on the spatial transcriptomics data. This included inspecting variable features and filtering out mitochondrial genes to clean the data for subsequent analysis. The single-cell RNA-seq data was prepared similarly, ensuring it was in an appropriate format for model input.

The Cell2location model was applied to deconvolute the spatial transcriptomics data, estimating cell type abundances at each spatial location. This involved integrating the spatial data with the reference single-cell RNA-seq data to create a comprehensive spatial map of cell distribution. Model parameters, fitting procedures, and diagnostic checks were carried out following best practices as recommended in Cell2location documentation and tutorials . The function cell2location.models.RegressionModel() prepares the AnnData object for compatibility with the Cell2location model, ensuring that the input data conforms to the expected format and structure. The function RegressionModel() initializes the RegressionModel object with the preprocessed and formatted AnnData. This step is crucial for specifying the model to be fitted to the spatial transcriptomics data. The rest of the parameters were the default model configurations.

Upon successful model convergence and estimation, the cell type abundance results were visualized using Matplotlib, with particular attention to the spatial distribution of different cell types within the liver sections. This step is critical for interpreting the cellular composition and architecture of the liver tissue, as well as understanding the heterogeneity and spatial organization of cell types.

## Competing interest statement

The authors declare the following competing financial interest(s): The Centrillion-affiliated authors are employees and the company may commercialize the work described herein.

9/26/24 7:27:00 PM

## Notes

### Summary of Updates

Additional sequencing data has been performed. That data is provided here, as well as a more thorough review of genes identified through unsupervised clustering.

